# Automated simulation-based membrane-protein refinement into cryo-EM data

**DOI:** 10.1101/2022.10.28.514175

**Authors:** Linnea Yvonnesdotter, Urška Rovšnik, Christian Blau, Marie Lycksell, Rebecca J. Howard, Erik Lindahl

**Affiliations:** Department of Biochemistry and Biophysics, Science for Life Laboratory, Stockholm University, Stockholm, Sweden; Department of Applied Physics, Science for Life Laboratory, KTH Royal Institute of Technology, Stockholm, Sweden

## Abstract

I.

The resolution revolution has increasingly enabled single-particle cryogenic electron microscopy (cryo-EM) reconstructions of previously inaccessible systems, including membrane proteins – a category that constitutes a disproportionate share of drug targets. We present a protocol for using density-guided molecular dynamics simulations to automatically refine atomistic models into membrane-protein cryo-EM maps. Using adaptive-force density-guided simulations as implemented in the GROMACS molecular dynamics package, we show how automated model refinement of a membrane protein is achieved without the need to manually tune the fitting force ad hoc. We also present selection criteria to choose the best fit model which balances stereochemistry and goodness-of-fit. The proposed protocol was used to refine models into a new cryo-EM density of the membrane protein maltoporin, either in a lipid bilayer or detergent micelle, and we found that results do not substantially differ from fitting in solution. Fitted structures satisfied classical model-quality metrics and improved the quality and the model-to-map correlation of the X-ray starting structure. Additionally, the density-guided fitting in combination with generalized orientation-dependent all-atom potential (GOAP) was used to correct the pixel-size estimation of the experimental cryo-EM density map. This work demonstrates the applicability of a straightforward automated approach to fitting membrane-protein cryo-EM densities. Such computational approaches promise to facilitate rapid refinement of proteins under different conditions or with various ligands present, including targets in the highly relevant superfamily of membrane proteins.

**STATEMENT OF SIGNIFICANCE:** Cryo-EM is an increasingly critical method of structure determination. As data collection and model generation become more efficient, iteratively fitting an experimental density can still require considerable time and expertise. Membrane proteins are particularly important targets in pharmacology and bioengineering, but can present distinctive challenges to data quality and modeling. Here, we tested a new tool to drive density fitting with molecular dynamics simulations, in context of a new structure of the membrane protein maltoporin. Fitting performed well in detergent, lipids, or solution, offering simpler options for fully automated simulation protocols. We were also able to apply fitting to adjust the microscope’s pixel size. The approach described here should be applicable to rapid, accurate refinement of a variety of membrane-protein structures.

## III. INTRODUCTION

Structure determination has been revolutionized by recent advances in cryogenic electron microscopy (cryo-EM). While the speed and ease of data acquisition have increased, the interpretation of the data by model building is increasingly the bottleneck. Manual model building and refinement are time consuming and require a high level of training and expertise. Automated approaches to *de novo* model building show promise to speed up the process [1–3]. However, there is still a need for additional refinement to improve the accuracy of models after building.

To improve the fit of a model to a density, model refinement may itself be automated by iteratively applying forces based on the experimental density to the model atoms. Additional forces, e.g. from dynamic elastic network models [4] or molecular dynamics [5, 6], aim to keep the model in a reasonable biochemical state. Recent approaches to overcoming challenges in such fitting include resolution Hamiltonian exchange, varying resolution [7], force constant replica exchange [8, 9], force fitting [10] and adaptive force-scaling [11]. Applications of automated fitting have accordingly expanded from low- to higher-resolution data, and from small to larger proteins [12, 13]. Open challenges remain in automating refinement of atomic models to cryo-EM data, particularly in the context of uncertain scaling factors and/or complex macromolecular targets.

Determining the structure of a membrane protein often involves a particular model-density discrepancy: detergents or lipids are required to solubilize the target, but give rise to cryo-EM densities that are highly variable, and may not directly correspond to physiological membranes. It is largely unclear how the cryo-EM density should be treated to appropriately model membrane regions in automated refinement. There are few examples available in the literature of automatic refinement of membrane proteins in a lipid environment. In work by Qi et al. [14], it is suggested that the simulation environment affects protein flexibility estimations derived from the refinement, while Mori et al. [15] report no effect on the result when fitting using an implicit micelle model [16] as compared to implicit solvent.

In this work we set out to determine the applicability of GROMACS density-guided simulations [11] to automatically refine a membrane protein system. Specifically, we collected and processed cryo-EM data for the *E. coli* membrane protein maltoporin, and used a previously reported X-ray structure as a starting model to refine against the cryo-EM density in various environments. In addition to identifying a robust parameter set and model-quality metrics for refinement of membrane proteins, we report an improved estimation of the pixel size of cryo-EM data based on goodness-of-fit and stereochemistry of simulated models.

## IV. MATERIALS AND METHODS

### A. Protein production and purification

Maltoporin is a protein endogenously expressed in *Escherichia coli* [17] *and it co-purified with a maltose binding protein fusion construct (a ligand-gated ion channel from Desulfofustis* deltaproteobacterium - DeCLIC [18]), which provided us with an interesting additional target to use for developing and assessing our refinement methodological work. The C43(DE3) *E. coli* cells were transformed with a pET-20b derived vector coding for DeCLIC-MBP and cultured overnight at 37°C. Cells were inoculated at 1:100 into 2xYT media with 300 *µ*g/ml ampicillin, grown at 37°C to OD_600_ = 0.4. After reaching the required OD_600_ the temperature was lowered to 20°C. The cells were further grown until the OD_600_ = 0.8, at which point they were induced with 100 *µ*M isopropyl-,*β*-D-1-thiogalactopyranoside (IPTG). Membranes were harvested from cell pellets that were ultracentrifuged in a supplemented buffer A (300 mM NaCl, 20 mM Tris-HCl pH 7.6, 1.5 ku benzonase nuclease, EDTA-free protease inhibitor cocktail). Harvested cell membranes were solubilized in 2% DDM, followed by amylose affinity purification and size exclusion chromatography to isolate the fusion protein and maltoporin.

### B. Cryo-EM sample preparation and data acquisition

Quantifoil 1.2/1.3 Cu 300 mesh grids (Quantifoil Micro Tools) were glow-discharged in the methanol vapor prior to sample application. 3 *µ*l sample was applied to the grid, which was then blotted for 3 s and plunge-frozen into liquid ethane using a FEI Vitrobot Mark IV. Data collection was carried out on an FEI Titan Krios 300 kV microscope with a post energy filter Gatan K2-Summit direct detector camera. Movies were collected at nominal 165,000x magnification, equivalent to a pixel spacing of 0.86Å. A total dose of 42 e^-^/Å^2^ was used to collect 40 frames over 6 sec, using a nominal defocus range covering −1.0 to −2.5 *µ*m.

### C. Image processing

All the image processing was carried out with the RELION 3.1 pipeline [19] (Figure S1), with its own implementation of MotionCorr used for motion correction. Defocus was estimated from the motion corrected micrographs using CtfFind4 [20]. Following manual picking, initial 2D classification was performed to generate references for autopicking. After autopicking particles were extracted, binned and an initial model with C3 symmetry was generated. Particles were then aligned by performing a 3D auto-refinement. The acquired alignment parameters were used to identify and remove noise through multiple rounds of pre-aligned 2D- and 3D-classification. Micelle density was eventually subtracted and the final 3D auto-refinement was performed using a soft mask covering the protein, followed by post-processing with the same mask. Local resolution was estimated using the RELION implementation. Per-particle CTF parameters were estimated from the resulting reconstruction using RELION 3.1. Global beam-tilt was estimated from the micrographs and correction applied. The final 3D reconstruction was generated, followed by post processing, producing a density used for fitting. After the pixel size was calibrated, the data was reprocessed from the beginning using RELION 4.0-beta-2 [21] at 0.83Å/px.

### D. Automated density-guided simulations

The previously determined maltoporin model (PDB ID: 1MAL) was used as a starting conformation [17] for automated refinement. The solution system was generated in GRO-MACS [22] using TIP3 water and 0.15 mM NaCl and energy minimized using the steepest descent algorithm that was allowed to run until convergence or a maximum force smaller than 10^*-*4^ kJ/mol. The CHARMM27 force-field was used for all GROMACS simulations [23, 24]. Detergent micelle embedding of 1MAL was performed using the CHARMM-GUI micelle builder [25] in 300 DDM molecules in a 175×175 nm (XY) box with 25 nm water thickness. The system was solvated with TIP3 water with a ion concentration of 0.15 mM NaCl. The system was energy minimized at 303.15 K within the micelle builder standard protocol. POPC membrane embedding of the initial model was performed using the CHARMM-GUI bilayer builder [25] in 733 POPC molecules and solvated in TIP3 water and 0.15 mM NaCl. The system was energy minimized at 303.15 K within the bilayer builder standard protocol.

Automated fitting into cryo-EM densities was performed by density-guided MD simulations using GROMACS 2021.3 [22]. Density-guided simulations were performed using the refined 3D density map as a bias. Alignment of the starting model to the density map was assessed and improved iteratively using VMD [26] and the GROMACS editconf functionality, rotating and translating the structure to ensure correct global alignment prior to starting the simulation [22]. The density-guided simulations were performed at 300K using adaptive force scaling starting at 10 kJ/mol with a feedback time-constant of 4 ps and a Gaussian transform spread-width of the model-generated density ranging between 0.68-0.731 Å (2*pixel size of target density map*0.425 as explained by Blau et al. [11]). Map similarities were calculated from their cross-correlation after normalization.

Evaluation of the overall fit of the model was performed every 2 ps using FSC average values with a threshold of 2.88 Å. FSC calculations were performed using a functionality available in a modified GROMACS version, available at https://gitlab.com/gromacs/gromacs/-/commits/fscavg. GOAP score source code was downloaded at https://sites.gatech.edu/cssb/goap/ [27]. GOAP scores were calculated for chain A every 2 ps and evaluated together with FSC average to find the best GOAP score within the FSC average plateau. The chosen best frame was energy minimized using the steepest descent algorithm that was allowed to run until convergence or a maximum force smaller than 10^*-*4^ kJ/mol/nm with the same long-range interaction settings as the density-guided molecular dynamics simulations. Phenix MolProbity scores were calculated for model quality assessment [28]. FSC-Q values were calculated using the Scipion3 implementation [29, 30] using default settings and a window size of 15. The generated FSC-Q maps are displayed at the default threshold as shown in ChimeraX [31].

### E. Pixel size calibration

An initial evaluation of the pixel size was done by analyzing the radius of gyration of the fit model normalized to the maltoporin starting model (PDB ID: 1MAL). Values for the radius of gyration were calculated using GROMACS[22]. Seven starting models were generated from an unrestrained simulation of maltoporin (PDB ID: 1MAL [17]). For each pixel size between 0.80 and 0.86 (0.80, 0.81, 0.82, 0.83, 0.84, 0.85 and 0.86), seven density-guided simulations were run using the cryo-EM data as target density. The GOAP score was calculated for chain A of the final frame of each simulation. The pixel size corresponding to the best average GOAP score was used as the correct pixel size estimation and the density map reprocessed accordingly.

### F. Data availability

Cryo-EM density maps of maltoporin in detergent micelles have been deposited in the Electron Microscopy Data Bank under accession number EMD-15903. The deposition includes the cryo-EM sharpened and unsharpened maps, both half-maps and the mask used for final FSC calculation, as well as the RELION FSC calculated output document [21]. Coordinates of the fitted model have been deposited in the Protein Data Bank, under the accession number 8B7V. The production runs, input files needed to generate a density-guided refinement and a list of commands are available via Zenodo https://doi.org/10.5281/zenodo.7257422.

## V. RESULTS

To build a membrane protein model in the context of a molecular-dynamics force field, it is unclear how extensive the model of the protein’s surrounding must be. Molecular-dynamics simulations typically benefit from describing the systems in question as completely as possible, including all particles and their interactions, within the computational resources available. However, for complex systems the setup itself is frequently non-trivial and can require extensive manual intervention, which becomes problematic in automated pipelines. Here, we first set out to test the influence of membrane mimetics of increasing complexity on density-guided simulations.

As a test system we used cryo-EM data for a native maltoporin sample purified from *E. coli*, solubilized in the detergent n-dodecyl-β-D-maltoside (DDM). The data were processed to 3.0 Å resolution (Figure S1, Table S3), and density corresponding to the detergent micelle was subtracted prior to final auto-refinement. As an initial model, we used an X-ray structure (PDB ID: 1MAL) previously reported at 3.1 Å [17]. To test the influence of membrane mimetics, we fit the cryo-EM density with initial model systems under three different conditions of increasing complexity. In one system, the model was solvated only by water, sodium and chloride ions, which can be performed fully automatically (Figure 1**A**, left). The second system had the protein embedded in a DDM micelle, using the density map prior to micelle subtraction to inform the number of detergent molecules needed to fill the experimental volume (Figure 1**A**, center). For the final system, a bilayer of 1-palmitoyl-2-oleoyl-sn-glycero-3-phosphocholine (POPC) lipid molecules was used as a simplified cell membrane (Figure 1**A**, right). Each model system was run with three replicates.

**FIG. 1.**
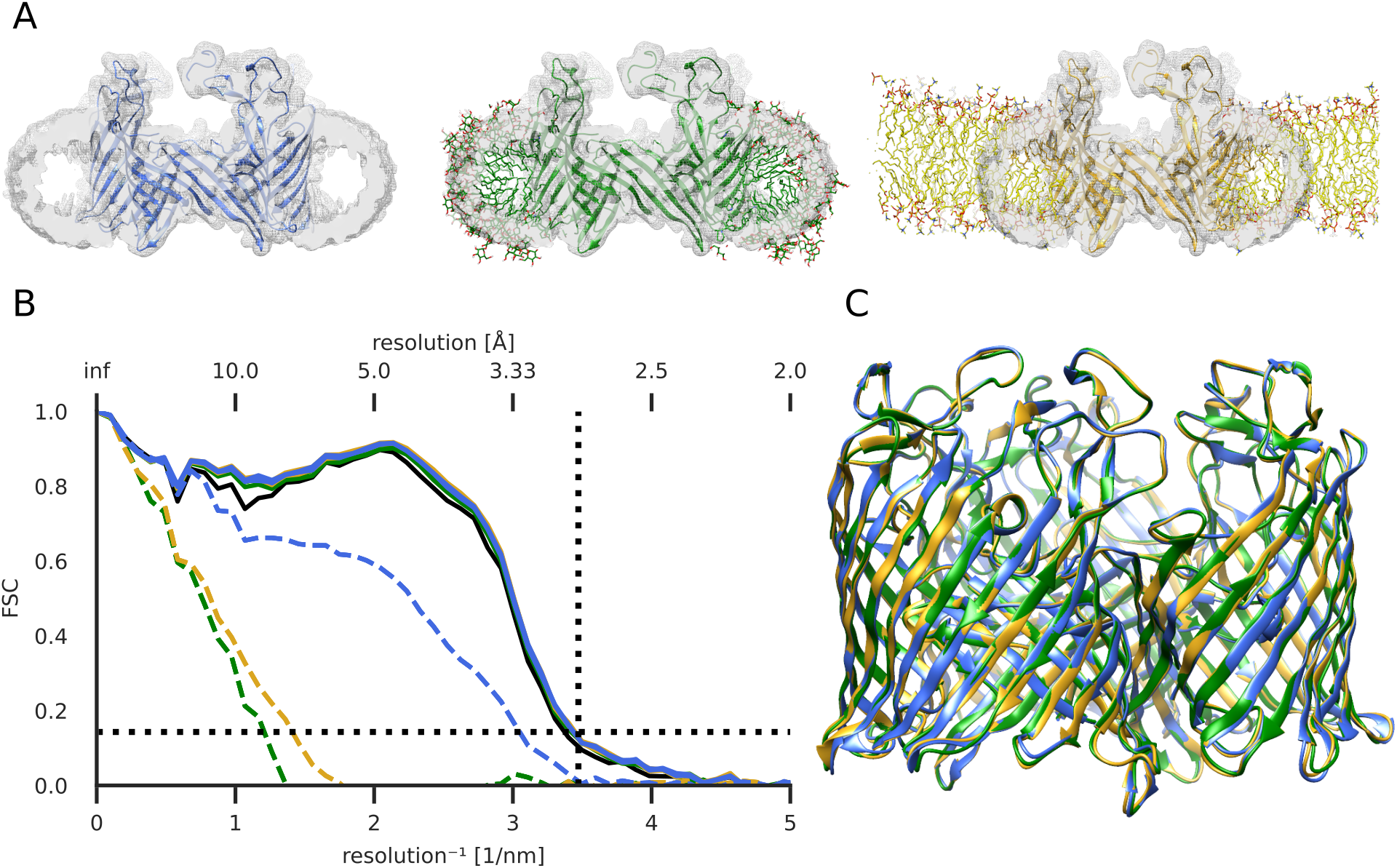
Effect of membrane mimetic on the quality of model fitting to detergent-solubilized maltoporin cryo-EM density. **A** - Cryo-EM density map (grey) prior to micelle subtraction, superposed with an initial model of maltoporin (PDB ID: 1MAL) in solution (left), DDM detergent micelle (middle) or POPC lipid bilayer (right). **B** - FSC of the best-fit model from density-guided simulations (n=3) of maltoporin in solution (blue), detergent (green) or lipid (yellow) fit into the same cryo-EM density map. FSC of initial models marked in dashed lines and Chimera fit-in-map rigid body fit model (solid black). Black dashed horizontal line at 0.143 and vertical at the lowest estimated local resolution of the map, 2.88 Å. **C** - Overlay of fitted models.

To minimize potential user bias from treating a manually built model as the ground truth, we monitored quality of fit of the refined model using Fourier shell correlation (FSC) to the target density (Figure S2**A-C**, dotted). All replicates of all systems achieved higher FSC to the density than did a fit-in-map rigid body fit of the starting structure performed in UCSF Chimera [32] (Figure S2**D-F**). The improvements were most notable in the nm size range, hinting at large scale conformational improvements.

From a molecular dynamics trajectory, the aim of density fitting is to extract a single model, which balances fit to the target density with stereochemical properties. Due to the increasing forces applied during the fitting process, inherent to adaptive force scaling [11], improvement in model-to-map agreement will at some point come at the cost of stereochemical deformations. We chose the best stereochemical model, as measured by generalized orientation-dependent all-atom potential (GOAP), a scoring function originally developed to assess protein structure prediction [27] (Figure S2**A-C**, solid). For each individual fitting simulation, the lowest (i.e. best) GOAP score corresponded to a point within the FSC_*average*_plateau, close to the maximum value. All simulations terminated within 3 ns due to excess adapted forces applied. Although models fit in solution (Figure S2**A**) achieved better GOAP scores than those in detergent (Figure S2**B**) or lipid (Figure S2**C**), the best-fit models were effectively superimposable (Figure 1**C**). Moreover, despite starting at different offsets compared to the target density map, there was no clear difference in best-fit model-to-map FSC between the three membrane-mimetic systems (Figure 1**B**, Figure S2**D-F**). As a final test of the stereochemical quality of our models and the influence of the membrane mimetic, we used MolProbity scores [28] and found improvement relative to the initial model in all conditions, with the best scores observed for models refined in solution (Table S1). The bestfit simulation frame from each replicate was energy-minimized with heavy-atom restraints, resulting in final models with even better MolProbity scores (Table S2).

To ensure that we made full use of the structural information while avoiding over-fitting, we further evaluated our structures in atomic detail using FSC-Q [29]. This quality measure compares model-to-map differences with differences between half-maps, and was recently integrated into the cryo-EM data processing framework Scipion3 [30]. Values of FSC-Q more negative than −0.5 indicate that an atom may have been refined to noise, while values above +0.5 indicate low correlation with the density or low resolution of the density. Best-fit models refined in solution, detergent or lipid had similar mean FSC-Q scores between 0.12 and 0.17 (Figure 2). Solution models had a higher percentage of atoms below −0.5 and above +0.5 than those in detergent or lipid; still, for all systems more than 93% of atoms had FSC-Q values in the well-fitted (−0.5 to +0.5) range.

**FIG. 2.**
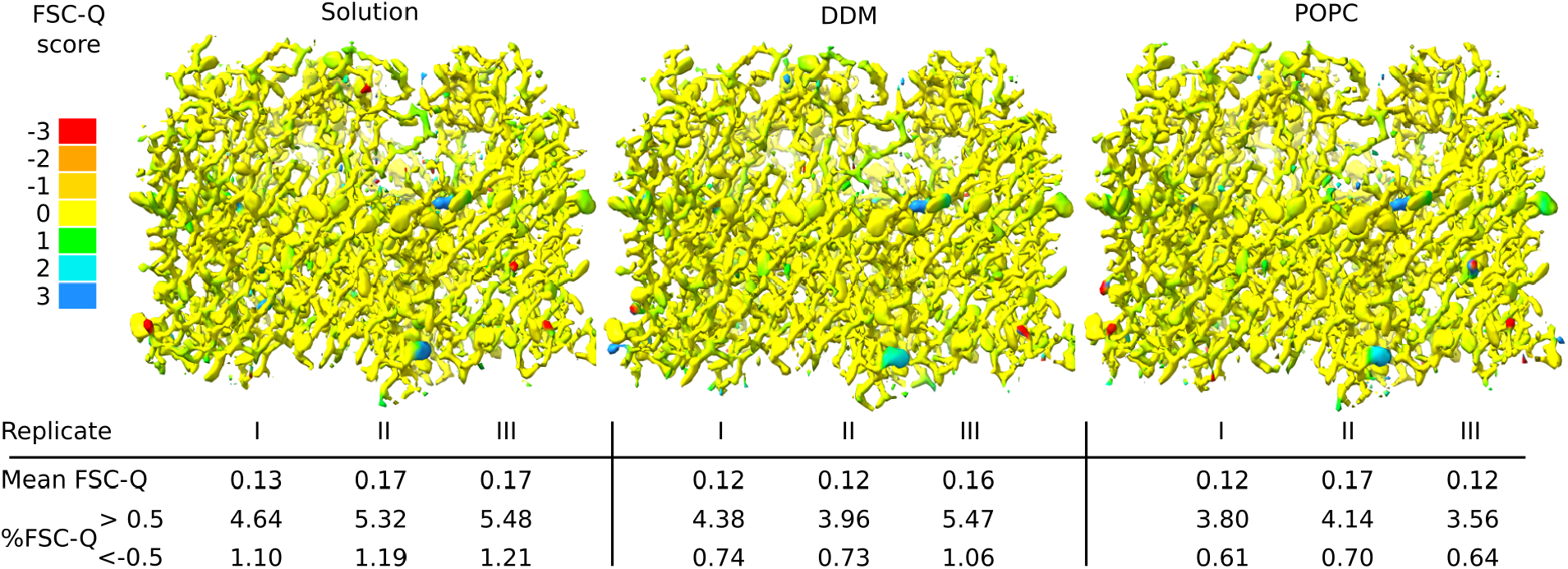
FSC-Q quality validation of models fit in solution (left), detergent (middle) or lipid (right). Densities generated from fit models, colored by residue FSC-Q values (>0.5 indicating poor correlation or low resolution of the density map, and <-0.5 indicating potential over-fitting). Mean FSC-Q values and percentage of atoms with values >0.5 and <-0.5 are indicated for three replicates (I-III) for each simulation set-up.

Interestingly, density-guided simulations provided an unanticipated opportunity to refine a parameter of our original cryo-EM reconstruction. Specifically, we sought to validate the pixel size estimate of our micrographs, reported at the time of data collection as 0.86 Å/pixel. For pixel size estimates that are too large, we expect an overall stretching of the model, while for too small estimates, we expect a compression. We refined the pixel size estimate with two approaches. First, we compared the overall shape of the model to a reference structure, the initial model (PDB ID: 1MAL [17]), during density-guided simulation, using radius of gyration. Due to the anisotropy of membrane systems, we calculated independently the radii of gyration around the membrane normal and two axes perpendicular to it, normalized to those of the reference structure. When fit to the original cryo-EM map processed at 0.86 Å/pixel, refinements converged to ∼3% larger radius of gyration than the reference structure (Figure 3**A**), suggesting that 0.86 Å/pixel may be larger than the actual pixel size during data collection.

**FIG. 3.**
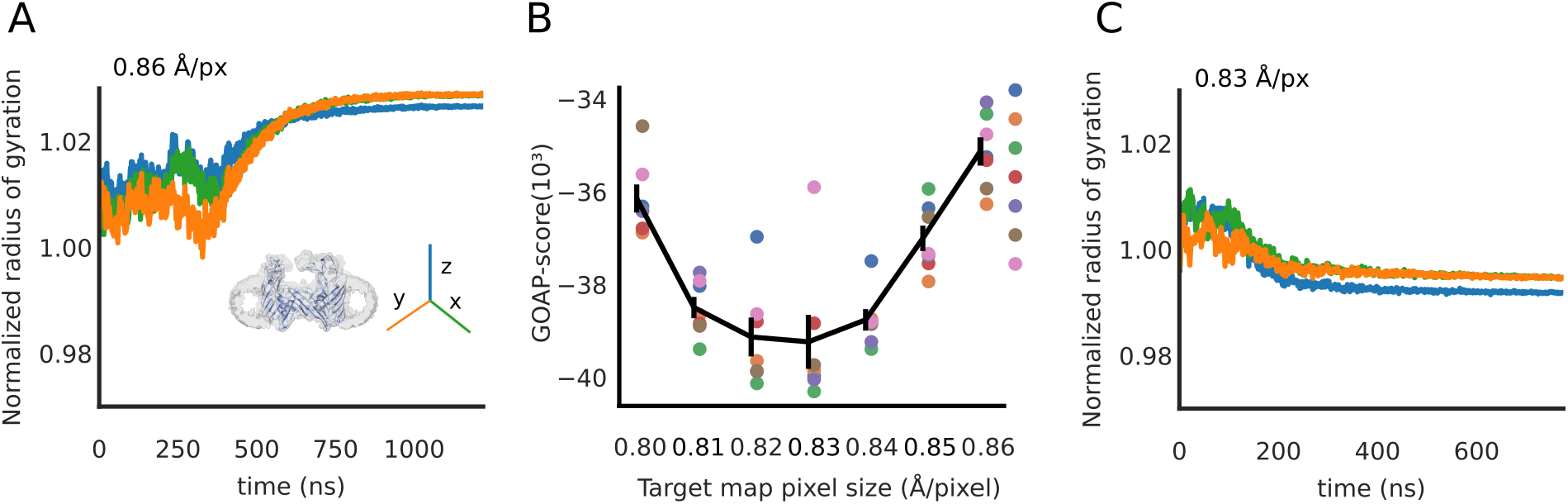
Pixel-size estimation improved by density-guided simulations. **A** - Radius of gyration of a fitted model, normalized to that of the initial model (PDB ID: 1MAL), guided by the experimental density map processed at the pixel size estimated at time of data collection (0.86 Å/pixel). Dimensions of the protein are plotted separately along x (green), y (orange) and z (blue) axes, the latter perpendicular to the membrane plane. **B** - Optimal GOAP scores for density-guided simulations initiated from a range of starting models (n=7, colored separately) guided by maps scaled at 0.80-0.86 Å/pixel. Solid line represents mean ± standard error of GOAP scores from fitting runs at each pixel size. **C** - Radius of gyration as in *A* for one model fitted to a map scaled at the optimized pixel size (0.83 Å/pixel).

To estimate the correct pixel size, we rescaled the cryo-EM map to voxel sizes ranging from 0.80 to 0.85 Å. Using a modified version of the approach suggested by Tiwari et al. [33], structures at seven different timepoints of an unrestrained molecular dynamics simulation of the initial model (PDB ID: 1MAL [17]) were used as seeds for density-guided refinement into the six re-scaled densities and the initial cryo-EM density (resulting in 7×7 = 49 fitting simulations). As any gross geometrical change is expected to deteriorate the overall stereo-chemical quality of the models, we used the average GOAP score to select the most suitable pixel size. This approach thus avoids reliance on a reference structure for the fitted model. Fitting to re-scaled maps improved the GOAP scores of best-fit models, with optimal scores at 0.83 Å/pixel (Figure 3**B**). The best-fit model refined at 0.83 Å/pixel converged to within 1 percent of the radius of gyration of the initial structure, again with relative contraction along the z-axis (Figure 3**C**). Subsequent discussions with the cryo-EM facility were consistent with a need to revise the pixel size estimate for our dataset, as well as others collected in that period of time. Accordingly, all analyses in this work (except Figure 3) correspond to cryo-EM data reprocessed from original micrographs at 0.83 Å/pixel.

## VI. DISCUSSION

Despite the recent rapid increase in cryo-EM structure depositions, membrane-protein structures remain underrepresented, particularly relative to their importance to physiological signaling and drug development. As data collection becomes increasingly efficient, automated modelling of atom coordinates into a three-dimensional map is poised to become a critical tool for interpreting cryo-EM data, and as a starting point for further molecular dynamics simulations. Here, we tested the applicability of a recently reported density-guided fitting tool [11] in the GROMACS software suite to membrane proteins and their membrane mimetics. We used new cryo-EM data for maltoporin, representative of a large class of beta-barrel transmembrane proteins [17, 34, 35]. We predicted that maltoporin’s relative rigidity would avoid large-scale conformational transitions confounding the fitting dynamics, and enable precise assessment of membrane-mimetic effects by minimizing restructuring of lipidic components around mobile hydrophobic regions.

As implemented here, the conditions chosen to mimic the embedding membrane – aqueous solution, detergent micelle or lipid bilayer – did not substantially influence the overall FSC between map and best-fit model. This agnosticism is consistent with previous indications of Mori et al., who achieved high-quality automated refinement using a different tool even in implicit solvent [15]. We attribute this seemingly surprising success of modelling a membrane protein in the absence of a solubilizing environment to several factors. To a lesser extent, we expect density-guided simulations to induce slightly hydrophobic behaviour by reducing side chain flexibility and thus the number of possible hydrogen bonding configurations [36]. We attribute a larger role to the forces from the density fitting, where an adaptive force constant adjusts the bias of the target density to overpower the influence of the embedding environment. This approach is similar to the replica exchange force constant in GENESIS [8], while different to the fixed force constant employed in MDFF [14].

The map-to-model quality measured by FSC-Q [29] was also comparable for all membrane-mimetic systems tested. We did however observe a slightly lower percentage of atoms with FSC-Q values below −0.5, indicating reduced over-fitting, in the detergent- and lipid-embedded models than in solution. We theorize that embedding in detergent or lipid might reduce fitting to noise left in a density map after micelle subtraction, by physically blocking that space. For density maps with high levels of noise, or with lipid or other density, not associated with the target protein, it might be advantageous to refine the protein in an embedded system; however, with well-defined density or tight mask, the difference should be small.

Models refined in solution achieved better GOAP scores than their detergent- or lipid-embedded counterparts. One contributing factor to this may be that GOAP is knowledge based score based on a set of 1011 non-homologous proteins with a resolution of <2Å available in the PDB [37]. As membrane proteins remain underrepresented in the PDB, GOAP score may represent the characteristics of proteins in solution better than a protein embedded in detergent or a lipid bilayer.

While most cryo-EM facilities perform regular pixel size calibrations to ensure correct scaling, there is sometimes a need for re-estimation. The change in radius of gyration to a reference structure has proven useful to give a first indication at the scale and direction of the error. A caveat to this approach is that it requires a known reference structure in the same conformational state as the target density. Further, taking an X-ray model as the ground truth for the overall shape of a protein is potentially problematic, as we lose the ability to find cryo-EM specific characteristics, which might stem from the difference in temperature at which structures are solved, or from the absence of crystal contacts. Instead, density-guided fitting in combination with GOAP scoring [27] can be used for such re-estimation, similar to the methodology proposed by Tiwari et al. [33]. This approach scores the soundness of the model stereochemistry, rather than comparing to a single reference. Again, as the GOAP score itself is based on a heuristic from a set of structures, mainly solved by X-ray crystallography [37], it might affect the results, as we compare between data derived from X-ray and cryo-EM. Finally, in order to deposit a final model to PDB it is necessary to re-process the data with the found correct pixel size, as no additional processing, such as re-scaling, of half-maps is allowed outside of the EM-processing tool for PDB deposition.

Adaptive force scaling allows sampling at different balances between stereochemistry and goodness of fit. Given that the cryo-EM density represents a thermodynamically favourable state, we expect that there is a corresponding minimum in the GOAP score of a correctly fit model to that density. In this case of maltoporin, devoid of large conformational changes, we find one GOAP score minimum at the FSC plateau. However, it should be possible to see two minima separated by a barrier when fitting between conformational states, one corresponding to the initial conformation and the other to the conformation represented by the target density. The here suggested protocol for automatically refining membrane proteins has been optimized for a relatively rigid protein. It is possible that some tweaking of the parameter set could aid in refining a more flexible protein. We then suggest increasing the time constant τ which regulates the interval at which the adaptive-force constant is increased or decreased, allowing for more sampling of the conformational space at each increment of the added bias. For large conformational changes we would expect an approach using density maps of increasing resolution, as reported by McGreevy et al. [38], combined with adaptive force scaling, to be a viable strategy. Here we show how density guided simulations can be used to refine protein models, however the same approach should be applicable to fitting lipids or ligands into experimental data.

## VII. CONCLUSION

In summary, we tested here an approach to automated model refinement of membrane proteins using density-guided simulations in GROMACS, consisting of the following steps: rigid-body alignment of an initial model to the density; setting up a solution-phase MD simulation; monitoring FSC_average_ and GOAP scores during density-guided simulations; selecting the simulation snapshot with the best GOAP score that falls on the FSC_average_ plateau; energy-minimizing the selected snapshot with heavy atoms restrained; and validating the final model on the basis of FSC, FSC-Q, GOAP score, and MolProbity score. Parallel simulations with variously scaled densities may also be useful in optimizing pixel-size estimation.

## VIII. ACKNOWLEDGMENTS

The authors would like to thank the Swedish Cryo-EM National Facility staff, especially Marta Carroni and Stefan Fleischmann from Stockholm and Michael Hall from Umeå, for kind assistance with data collection. The Facility is funded by the Knut and Alice Wallenberg Foundation, Erling Persson and Kempe Foundations. This project was supported by the Swedish Research Council (2019-02433, 202105806), the Swedish e-Science Research Centre, and the BioExcel Center of Excellence (EU-823830). Computational resources were provided by the Swedish National Infrastructure for Computing (SNIC).

## IX. AUTHOR CONTRIBUTIONS

Conceptualisation: LY, UR, CB; methodology: LY, CB; software: LY, CB, UR; validation: LY; formal analysis: LY, UR, ML, CB, RH, EL; investigation: LY, UR, CB, ML; data curation: LY, UR; original draft: LY; review and editing: LY, UR, ML, CB, RH, EL; visualization: LY, UR; supervision: RJH, EL; project administration: RJH, EL; funding acquisition: EL.

## XII. TABLES

**TABLE 1.**
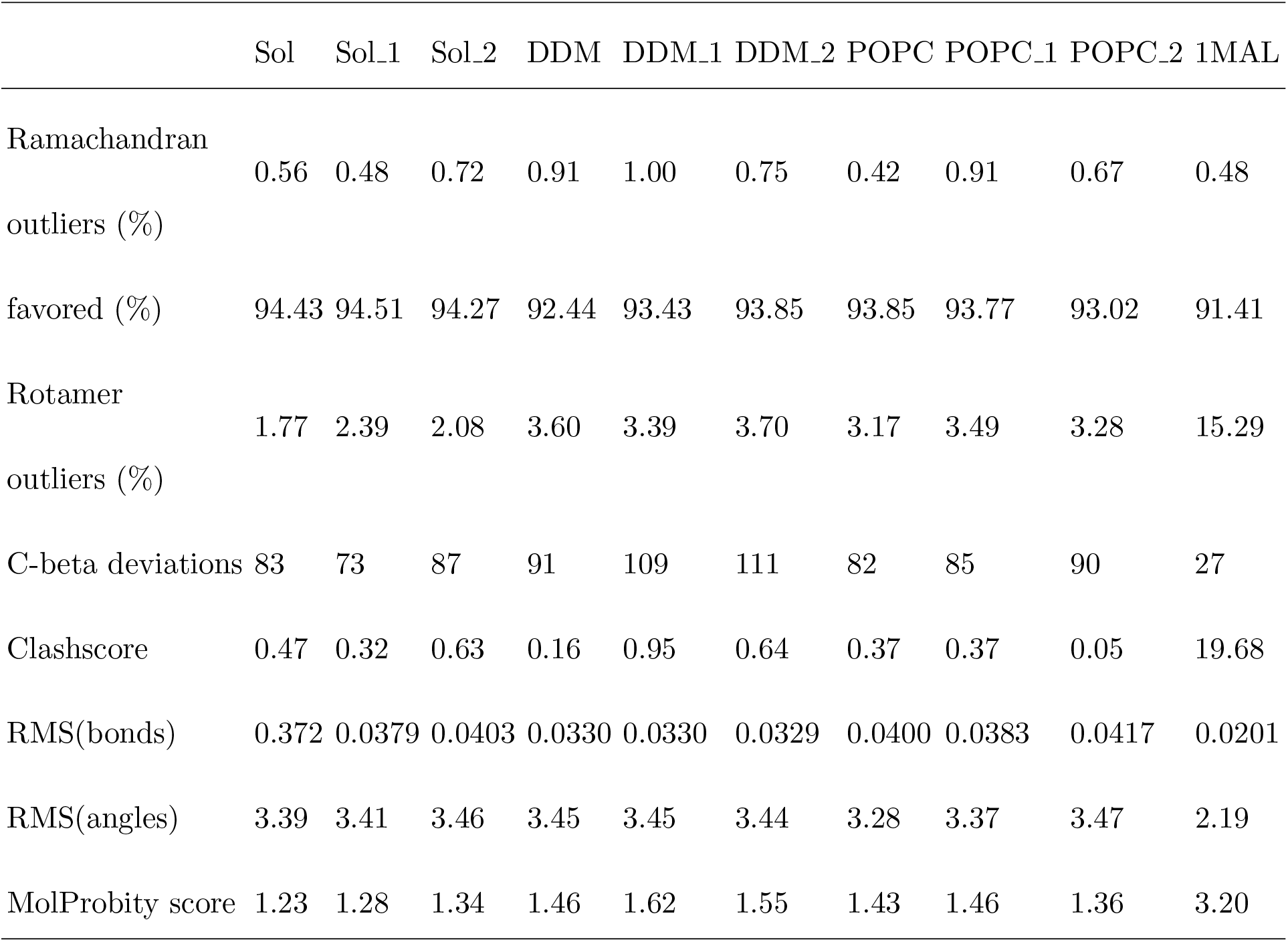
Model quality statistics at the best fit frame of the density guided simulation trajectories (n=3) in solution (Sol, Sol_1, Sol_2), DDM (DDM, DDM_1, DDM_2) and POPC (POPC, POPC_1, POPC_2).

**TABLE 2.**
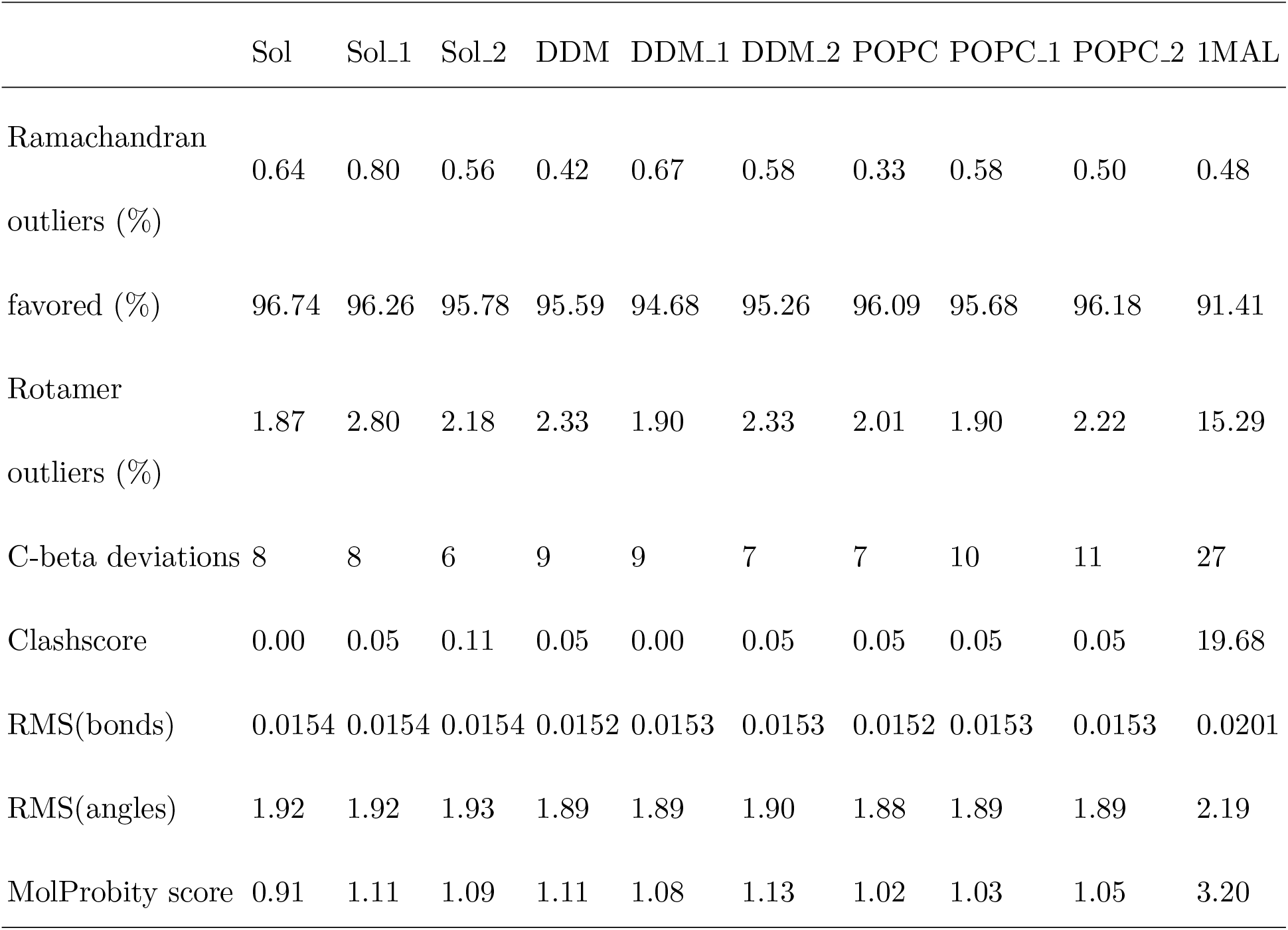
Model quality statistics of best fit frame of the density guided simulation trajectories (n=3) after heavy atom restrained energy minimization (Sol, Sol_1, Sol_2), POPC (POPC, POPC_1, POPC_2) and DDM (DDM, DDM_1, DDM_2).

**TABLE 3.** Cryo-EM data collection statistics

|  |  |
| --- | --- |
| *Data Collection* |  |
| Microscope | FEI Titan Krios |
| Magnification | 165,000 |
| Voltage (kV) | 300 |
| Electron exposure ($e^-/\text{\AA}^2$) | $\sim 40$ |
| Defocus Range ($\mu\text{m}$) | -2.0 – -3.6 |
| Pixel Size ($\text{\AA}$) | 0.83 |
| Symmetry Imposed | C3 |
| Number of Images | 3740 |
| Particles Picked | $\sim 579,000$ |
| Particles Refined | 37,521 |
| *Refinement* |  |
| Resolution ($\text{\AA}$) | 3.0 |
| FSC Threshold | 0.143 |
| Map sharpening B-factor | -70 |

