## Supplementary Information for "Automated simulation-based membrane-protein refinement into cryo-EM data"

### XIII. SUPPLEMENTARY INFORMATION

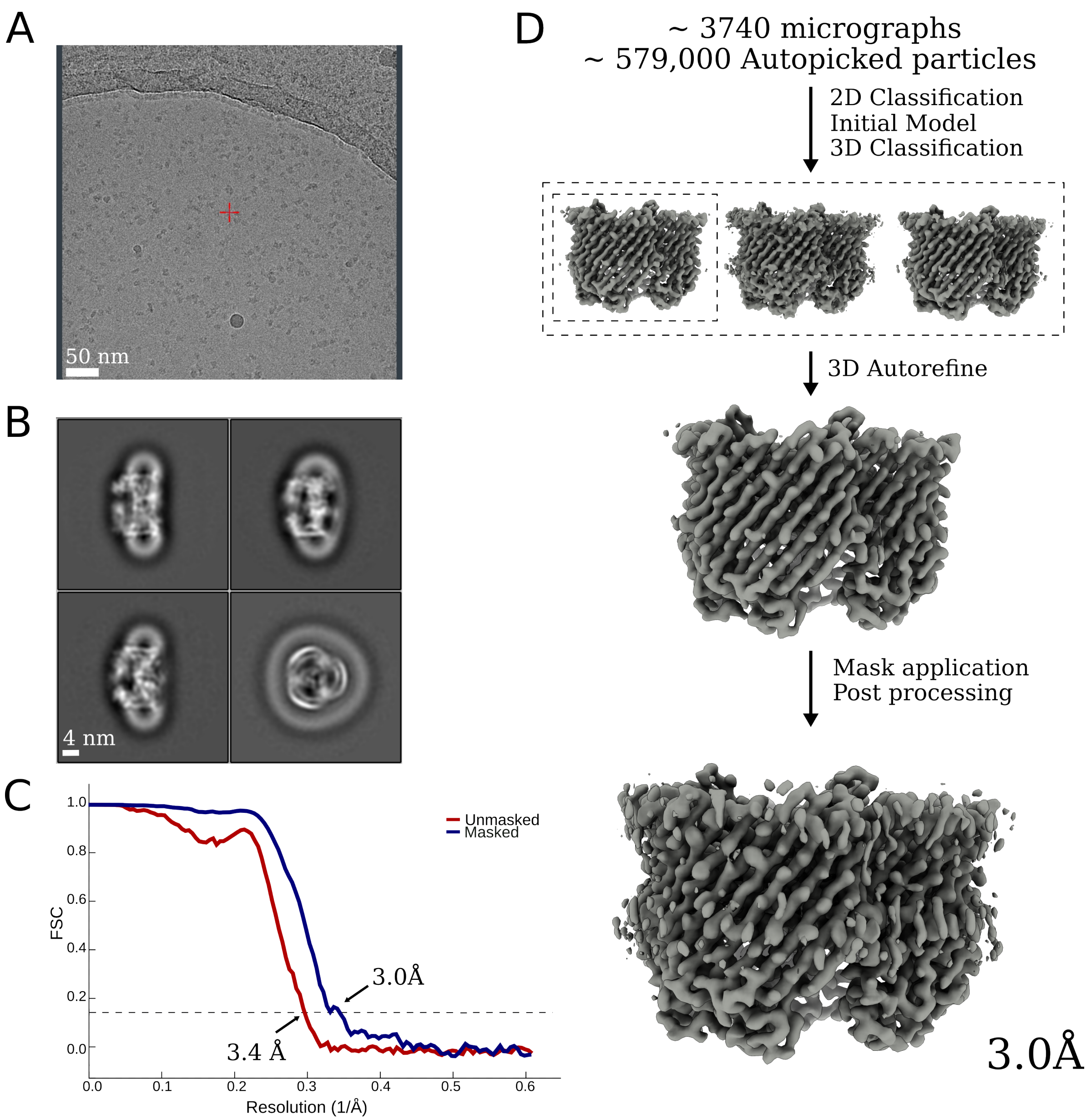


FIG. S1. Maltoporin cryo-EM processing pipeline. **A** - Cryo-EM representative micrograph from the data collection. **B** - Representative 2D classes in multiple different orientations. **C** - FSC curves for unmasked (*red*) and masked (*blue*) map. **D** - Maltoporin processing pipeline with representative 3D classes, 3D refined density and a sharpened final map.

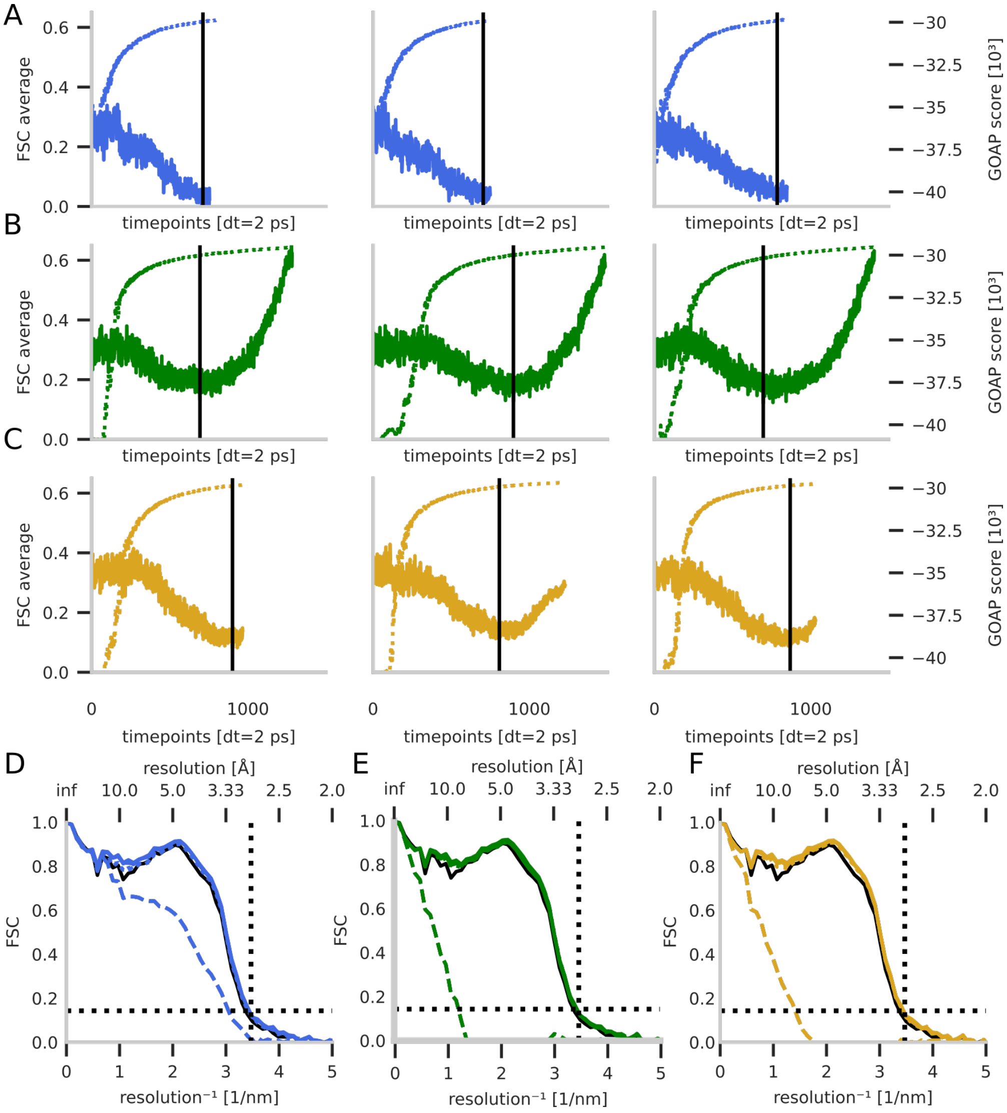


FIG. S2. **A-C** - FSC$_{average}$ (dotted) and GOAP score (solid) over the refinement simulations (n=3) in solution (A), DDM (B) and POPC (C), best GOAP score frame marked with vertical black line. **D-F** - FSC of best fit position of density-guided simulations (n=3) of maltoporin (1MAL) in solution (blue), in DDM detergent micelle (green) and embedded in POPC (yellow) fit into cryo-EM density map, energy minimized models (dotted colored lines). Respective starting position marked in dashed lines and chimera fit-to-map rigid body fit starting model (solid black). Black dotted horizontal line at 0.143 and vertical at the lowest estimated local resolution of the map, 2.88 Å.
